# Meiotic Genes in Colpodean Ciliates Support Secretive Sexuality

**DOI:** 10.1101/132472

**Authors:** Micah Dunthorn, Rebecca A. Zufall, Jingyun Chi, Konrad Paszkiewicz, Karen Moore, Frédéric Mahé

## Abstract

Colpodean ciliates potentially pose a problem to macro-organismic theories of evolution: they are putatively asexual and extremely ancient, and yet there is one apparently derived sexual species. If macro-organismic theories of evolution also broadly apply to microbial eukaryotes, though, then most or all of the colpodean ciliates should merely be secretively sexual. Here we show using de novo genome sequencing, that colpodean ciliates have the meiotic genes required for sex and these genes are under functional constraint. Along with these genomic data, we argue that these ciliates are sexual given the cytological observations of both micronuclei and macronuclei within their cells, and the behavioral observations of brief fusions as if the cells were mating. The challenge that colpodean ciliates pose is therefore not to evolutionary theory, but to our ability to induce microbial eukaryotic sex in the laboratory.

## INTRODUCTION

There are many costs to meiotic sex (Lehtonen *et al.*, 2012; Maynard Smith, 1978). Sex is nevertheless maintained in animals and plants because it allows, for example, quicker escapes from parasites and quicker adaptations to changing environments (Bell, 1982; Hamilton, 2001; Maynard Smith, 1978). As asexuality is thus thought to lead to extinction, putative asexual macro-organisms are intensely studied (Normark *et al.*, 2003). Even in light of this macro-organismic theory, many microbial eukaryotic species and higher clades have often been considered to be asexual (Fenchel and Finlay, 2006; Schlegel and Meisterfeld, 2003; Sonneborn, 1957). One of the largest putative asexual microbial eukaryotic groups are the Colpodea ciliates (Foissner, 1993).

Ciliates have dimorphic nuclei within each cell: micronuclei, which are transcriptionally inactive during vegetative growth; and macronuclei, which produce all mRNA required for protein synthesis (Lynn, 2008). Ciliate sex—called conjugation—occurs by brief cell fusion of complementary mating types and mutual exchange of haploid products of micronuclear meiotic division (Bell, 1988; Phadke and Zufall, 2009; Zufall, 2016). Although sex has been widely observed in almost all ciliate groups, some ciliates are asexual because they lack micronuclei (Zufall, 2016). Many of these amicronucleate strains are closely related to known sexual ciliates, but some are potentially old (Doerder, 2014).

The putative asexual colpodean ciliates form a clade of over 200 described species that all have micronuclei and macronuclei. They have baroque morphologies in their somatic and oral regions, and are primarily found in terrestrial environments (Foissner, 1993; Lynn, 2008). Colpodeans have been consistent in their lack of conjugation in the laboratory even after more than forty years of observations, with the exception of only one species: *Bursaria truncatella* (Foissner (1993); Figure 1). Pseudoconjugation—where cells from clonal lines briefly fuse as if to mate—has been observed in other colpodeans while living in petri dishes, but there is no apparent exchange of haploid meiotic products (Foissner, 1993).

**Fig. 1.**
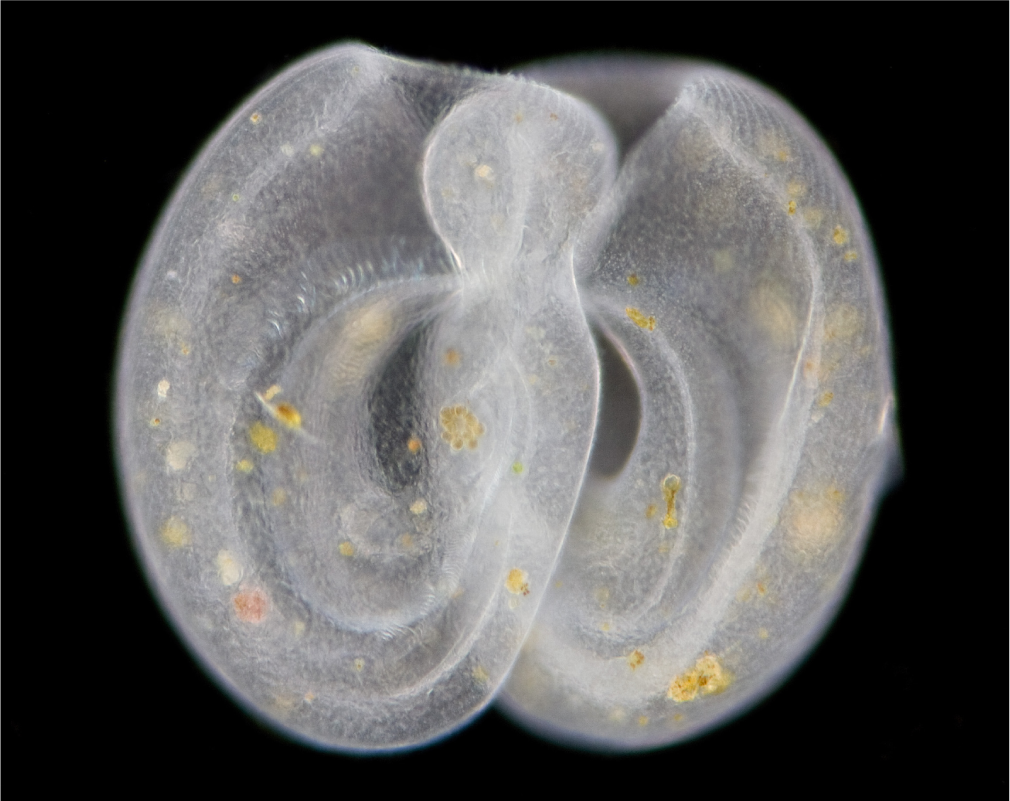
Two *Bursaria truncatella* cells during conjugation (= ciliate sex). Out of over 200 described species of colpodean ciliates, only *B. truncatella* has directly been observed to have sex. Copyright © Charles Krebs.

Because of macro-organismic theory about the maintenance of sex, two lines of evolutionary theory were previously used to argue that all, or almost all, of the colpodeans are secretively sexual (Dunthorn and Katz, 2010). First, the colpodeans may have originated before the Phanerozoic (Wright and Lynn, 1997), but ancient asexual lineages are extremely rare (Martens *et al.*, 2003; Maynard Smith, 1978; Normark *et al.*, 2003). Second, *B. truncatella* is in a phylogenetically derived position (Dunthorn *et al.*, 2011, 2008) implying a loss and then regain of sex, but reversing the loss of complex traits is thought unlikely to occur (Collin and Miglietta, 2008; Gould, 1970; Teotónio and Rose, 2001).

In the absence of direct observations of sex in the Colpodea, the most powerful approach to evaluate secretive sexuality in this ciliate clade is to look for meiotic genes, as meiosis is the central aspect of eukaryotic sex (e.g., Chi *et al.* (2014a); Malik *et al.* (2008); Ramesh *et al.* (2005); Schurko and Logsdon (2008)). We therefore sequenced the genomes of the known sexual colpodean *B. truncatella* and the putative asexual *Colpoda magna*. With these genomic data, we inventoried meiotic genes to evaluate their presence or absence, and we evaluated the rate of evolution of the inventoried genes relative to the same genes in known sexual ciliates from other clades (Chi *et al.*, 2014b) to look for evidence of relaxed selection.

## MEIOTIC GENE INVENTORY AND EVOLUTIONARY RATES

To uncover the genes involved in meiosis in *B. truncatella* and *C. magna*, clonal cell lines were *de novo* genome sequenced. The aim of the sequencing was to produce open reading frames, not to resolve issues of chromosomal scaffolding or differences between micronuclei and macronuclei. From these open reading frames, we evaluated the presence or absence of 11 meiosis-specific genes, and 40 meiosis-related genes that are also involved in mitosis.

The complement of meiotic genes in the two colpodeans generally matched those from sexual ciliates (Fig. 2). Both *B. truncatella* and *C. magna* had *SPO11*, which causes the double-strand DNA breaks that initiate meiosis (Keeney, 2001). Six other meiosis-specific genes that are involved in crossover regulation were uncovered: DMC1, HOP2 (but not in *C. magna*), *MER3* (which was not found in other ciliates), *MND1*, *MSH4*, and *MSH5*. Like other ciliates, *HOP1*, *RED1*, and *ZIP1* were not found in the colpodeans, supporting the view of Chi *et al.* (2014b) that ciliates in general have a slimmed crossover pathway 1 that lacks a synaptonemal complex. *B. truncatella* and *C. magna* mostly had the same complement of meiosis-related genes as found in the other ciliates, including *MUS81*; except that *CDC2*, *MPH1/FANCM*, *MPS3/SUN-1/SAD1*, *SGS1*, and *SLX1* were missing in one or both of them.

**Fig. 2.**
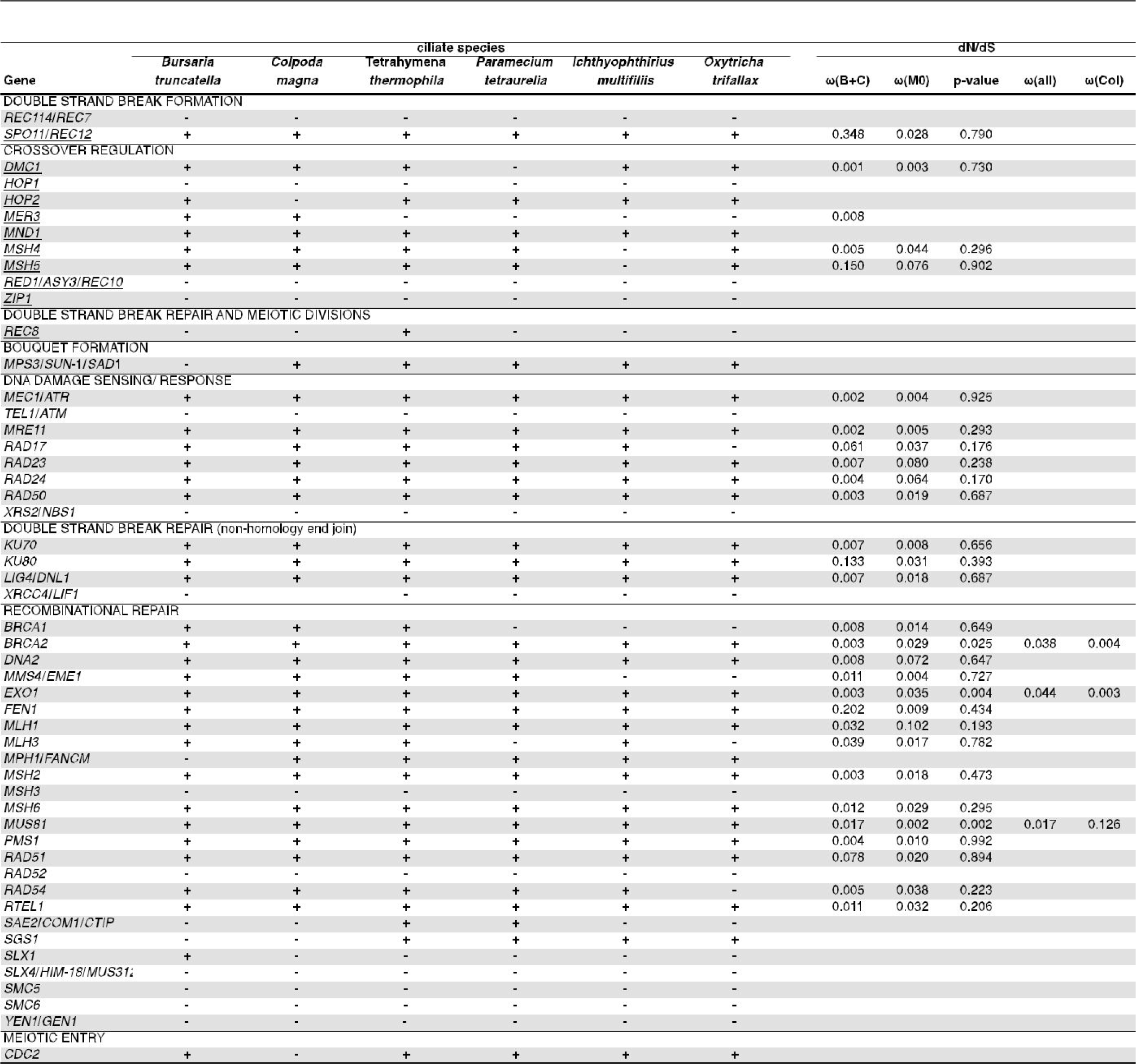
Meiosis genes inventoried in two colpodean ciliates: *Bursaria truncatella* and *Colpoda magna*. Genes are grouped according to functions. Meiosis-specific genes are underlined. Data from *Tetrahymena*, *Paramecium*, *Ichthyophthirius*, and *Oxytricha* are from (Chi *et al.*, 2014b). ω(B+C) indicates the value of dN/dS between *B. truncatella* and *C. magna*. ω(M0) indicates the value of dN/dS for all taxa. The significance of the difference between a model with one parameter for ω vs. a model with an addition parameter for ω on the *C. magna* branch is shown by the p-value from a *χ*^2^ test with one degree of freedom. In cases where this test is significant, ω(all) indicates the background rate and ω(Col) is the rate for *C. magna*.

To look for evidence of relaxed selection in the uncovered meiotic genes from the colpodeans, we evaluated the rates of nonsynonymous relative to synonymous substitutions (ω; Fig. 2). Genes showing evidence of relaxed selection (that is, elevated ω) would indicate the loss of functional constraints due to loss of use in sexual reproduction (Lahti *et al.*, 2009). Evolutionary rates were first measured in the genes contained in both *B. truncatella* and *C. magna*. In this pairwise comparison, all estimated values of ω were much less than one, indicating strong purifying selection in the meiotic genes. A second comparison was made of the two colpodeans with homologs from other ciliate species, where estimated values of ω were also very low. For this comparison between all of the colpodeans and other ciliates, two models were evaluated: M0, where all species were modeled to evolve at the same rate; and M2, where all evolve at the same rate except *C. magna*. If the genes in *C. magna* were experiencing relaxed selection, in contrast to purifying selection in the other taxa, we would expect to find evidence of larger values of ω in the *C. magna* lineage. However, only three genes show a significant difference in evolutionary rates in *C. magna* (that is, the p-value for the log-likehood ratio test was ≤ 0.05), and of those three only one, *MUS81*, is in the predicted direction.

## GENOMIC DATA SUPPORT SECRETIVE SEX IN COLPODEANS

The meiotic genes found in this inventory of the two colpodeans were the same as those found in known sexual ciliates. These meiotic genes would have been lost in *C. magna* if the colpodeans are asexual. In addition, there was no evidence of relaxed selection on these genes in *C. magna*, suggesting that these genes are under functional constraint just as in the sexual species. As predicted (Dunthorn and Katz, 2010), the colpodean ciliates are likely sexual given these genomic data.

It should be noted, however, that having functional meiotic genes can also allow for activities other than what we defined here as sex (where there is a requirement of recombination between two individuals). These meiotic genes could be used in canonical genetic pathways such as automixis (= selfing), which could allow for the purging of deleterious alleles, but would not provide the benefit of genetic exchange between individuals. They can also be used in non-canonical genetic pathways such as: diplomixis in *Giardia intestinalis*, where there is homologous recombination between the two nuclei within each cell but no meiotic reduction in ploidy (Carpenter *et al.*, 2012; Poxleitner *et al.*, 2008); parasexuality in *Candida albicans*, where tetraploidy caused by cell fusion is non-meiotically reduced to diploidy (Bennett and Johnson, 2003; Forche *et al.*, 2008); and DNA repair in bedelloid rotifers from damages induced by desiccation and UV radiation (Bininda-Emond *et al.*, 2016; Gladyshev and Meselson, 2008; Hespeels *et al.*, 2014).

Beyond these genomic data, there are two additional lines of support for sexuality in colpodean ciliates. One is cytological. The other is behavioral.

All colpodean species that have been described in enough detail have micronuclei (e.g., Bourland *et al.*, 2013; Dunthorn *et al.*, 2009; Foissner, 1993; Foissner *et al.*, 2014; Quintela-Alonso *et al.*, 2011). As micronuclei are only involved in sex (as far as we know), we propose that this cytological feature would have been lost over evolutionary time if the colpodeans are asexual. While the colpodeans have micronuclei, at least six species from two different subclades have micronuclei and macronuclei with shared outer nuclear membranes (Dunthorn *et al.*, 2008; Foissner, 1993). This shared outer membrane could possibly chain the micronucleus to the macronucleus, and thereby prevent it from participating in sex. That is, having micronuclei chained to macronuclei could be analogous to having no micronuclei at all. However, it is unknown if this shared outer-nuclear membrane is present in most individuals within those six species or if the micronuclei can break free at some point during the cell cycle.

Pseudoconjugation has been observed in some colpodean ciliates while living in petri dishes (Foissner, 1993). As conjugation is only involved in sex (as far as we know), we propose that this behavioral feature would have been lost over evolutionary time if the colpodeans are asexual. It should be noted, however, that pseudoconjugation can also allow for activities other than what we defined here as sex. For example, pseudocopulation occurs in the all female *Aspidoscelis uniparens* (desert grassland whiptail lizards), where females need to be mounted by other females to induce parthenogenesis (Crews and Fitzgerald, 1980).

## CONCLUSIONS

Our genomic analyses show that ciliates do no violate the macro-organismic theories against ancient asexuals and the loss-and-regain of complex characters: the Colpodea are sexual. This finding supports the increasingly accepted view that sex in in microbial eukaryotes is ubiquitous although often secretive (Dunthorn and Katz, 2010; Speijer *et al.*, 2015). Such secretive sex may result in long periods of mitotic division without meiosis, which could lead to the buildup of high mutational loads in the quiescent germline genomes of the colpodeans. However, rare sex could be tolerated if the colpodeans have extremely low base-substitution mutation rates as found in the ciliates *Paramecium tetraurelia* and *Tetrahymena thermophila* (Long *et al.*, 2016; Sung *et al.*, 2012).

## MATERIAL AND METHODS

There are many ways to define sex (Lehtonen and Kokko, 2014). Here we use the definition by Normark *et al.* (2003): “meiosis followed by the fusion of meiotic products from different individuals.” This definition excludes both recombination in bacteria and automyxis by eukaryotes.

Cells of *B. truncatella* were obtained from Carolina Biological Supply Company (Burlington, NC, U.S.A.) and cells of *C. magna* were obtained from the American Type Culture Collection (#50128). Clonal cultures were established and grown in Volvic water with wheat grains and *Klebsiella*. *Bursaria truncatella* cultures also included *Paramecium*. Individual cells were picked with a pipette and washed three times in sterilized Volvic water, then allowed to starve for 48 hours. Ten starved cells from each species were individually whole genome amplified with REPLI-g Mini Kit (Hilden, Germany) following manufacturer’s instructions. For each species, the ten whole-genome amplified products were combined in equal DNA concentrations.

Amplified DNA from *B. truncatella* was sequenced with Illumina MiSeq v2 chemistry (13,288,644 2x250 bp reads) and Illumina HiSeq v3 chemistry (113,546,269 2x150 bp reads). Amplified DNA from *C. magna* was sequenced with MiSeq v3 chemistry (18,770,554 2x300 bp reads) and HiSeq v3 chemistry (28,298,554 2x150 bp reads). The optimal k-mer length for genome assembly was searched within a 21-201 range with KmerGenie v1.6976 (Chikhi and Medvedev, 2014), using the “diploid” parameter. Genomes were then assembled with Minia v2.0.7 (Chikhi and Rizk, 2012), setting the kmer minimal abundance to 5. The obtained contigs were then analyzed with AUGUSTUS v2.7 (Stanke *et al.*, 2004) for a structural annotation, using the following parameters: search on both strands; genome is partial; predict genes independently on each strand, allow overlapping genes on opposite strands; report transcripts with in-frame stop codons; species set to the ciliate *T. thermophila*. Reads were deposited in GenBank’s Sequence Read Archive under BioProject numbers PRJNA381863 and PRJNA382551.

A query database of 11 meiosis-specific genes and 40 meiosis-related genes from ciliate and non-ciliate eukaryotes was established using literature and keyword searches of the NCBI protein database was taken from Chi *et al.* (2014b). For *REC8*, we used canonical eukaryotic sequences and *T. thermophila*’s non-canonical *REC8* (Howard-Till *et al.*, 2013). The ORFs of the two colpodeans were searched by the query database using BlastP (Altschul *et al.*, 1990) and HMMER v3.0 (Eddy, 2001). Hits with E-values *<* 10^-4^ for the full sequence were retained. Verification of candidate homologs used reciprocal BlastP search against the non-redundant protein sequence database of NCBI (Supplementary File 1).

In order to determine the strength of purifying selection acting on the inventoried meiotic genes, we measured ω = dN/dS, the number of nonsynonymous substitutions per nonsynonymous site divided by the number of synonymous substitutions per synonymous site. Sequences generated from this study were aligned with homologous sequences identified from *T. thermophila*, *P. tetraurelia*, *Ichthyophthirius multifiliis*, and *Oxytricha trifallax* by (Chi *et al.*, 2014b). Sequences were aligned in Geneious v4.8.3 (Kearse *et al.*, 2012) using Translation Align with ClustalW v2 (Larkin *et al.*, 2007). Maximum Likelihood genealogies of each gene were inferred with MEGA7 (Kumar *et al.*, 2016), and synonymous and nonsynonymous substation rates were estimated with PAML v4.8 (Yang, 2007). ω was first calculated between *B. truncatella* and *C. magna*. Then all species were included to test whether the lineage leading to *C. magna* exhibited higher values of ω, which would be expected if these genes were no longer functional and thus experiencing relaxed selection. Using codeml, we compared a model of evolution with one value of ω for the whole tree (model = 0) to a model where the *C. magna* lineage had a separate value of ω (model = 2). A log-likelihood ratio test was used to determine whether the second model provided a significantly better fit to the data.

## ACKNOWLEDGEMENT

This work was supported by the: Deutsche Forschungsgemeinschaft [grant DU1319/1-1] to M.D.; National Institutes of Health [grant R01GM101352] and the University of Houston [GEAR] to R.A.Z.; and Wellcome Trust Institutional Strategic Support Fund [grant WT097835MF], Wellcome Trust Multi User Equipment Award [grant WT101650MA], and Medical Research Council Clinical Infrastructure Funding [grant MR/M008924/1] to K.P. and K.M.

